# Plant growth and phytoactive alkaloid synthesis in *Mitragyna speciosa* (kratom) in response to varying radiance

**DOI:** 10.1101/2021.10.18.464895

**Authors:** Mengzi Zhang, Abhisheak Sharma, Francisco León, Bonnie Avery, Roger Kjelgren, Christopher R. McCurdy, Brian J. Pearson

## Abstract

The dose-dependent consumptive effect of kratom and its potential application as an alternative source of medicine to mitigate opioid withdrawal symptoms has brought considerable attention to this plant. Increased interest in the application and use of kratom has emerged globally, including North America. Although the chemistry and pharmacology of major kratom alkaloids, mitragynine and 7-hydroxymitragynine, are well documented, foundational information on the impact of plant production environment on growth and kratom alkaloids synthesis is unavailable. To directly address this need, kratom plant growth, leaf chlorophyll content, and alkaloid concentration were evaluated under three lighting conditions: outdoor full sun, greenhouse unshaded, and greenhouse shaded. Nine kratom alkaloids were quantified using an ultra-performance liquid chromatography-tandem mass spectrometry (UPLC-MS/MS) method. Contents of six alkaloids to include: mitragynine, speciogynine, speciociliatine, mitraphylline, coynantheidine, and isocorynantheidine were not significantly impacted by lighting conditions, whereas 7-hydroxymitragynine was below the lower limit of quantification across all treatments. However, paynantheine concentration per leaf dry mass was increased by 40% and corynoxine was increased by 111% when grown under shade conditions in a greenhouse compared to outdoor full sun. Additionally, total alkaloid yield per plant was maximized when plants were under such conditions. Greenhouse cultivation generally promoted height and width extension, leaf number, leaf area, average leaf size, and total leaf dry mass, compared to outdoor full sun condition. Rapid, non-destructive chlorophyll evaluation correlated well (r^2^ = 0.68) with extracted chlorophyll concentrations. Given these findings, production efforts where low-light conditions can be implemented are likely to maximize plant biomass and total leaf alkaloid production.

## Introduction

*Mitragyna speciosa*, commonly known as kratom, is a tropical small to medium size (4-16 m) tree indigenous to wetland forests of Southeast Asia. Historically, kratom was used in Thailand, Malaysia, and Indonesia to serve as a mild herbal stimulant, pain reliever, and to treat diarrhea and opium addiction [1–3]. In southeast Asia, kratom leaves are harvested and consumed fresh by chewing or steeping in water to make tea [3]. In the Western hemisphere where fresh kratom is unavailable, kratom is sold in the form of dried and ground powder or as a concentrated liquid extract for easier transportation and consumption [4].

Kratom produces an array of psychoactive compounds. So far more than 54 compounds including alkaloids, flavonoids, and terpenoids have been identified within kratom [5–7]. Although kratom’s alkaloids are likely produced by the plant to aid in defense of environmental challenges, they have demonstrated activity at central nervous system targets and may be medically valuable for human health [8–10]. Of the wide array of alkaloids found in kratom leaves, mitragynine and 7-hydroxymitragynine are the best understood and considered the most psychoactive [6]. Mitragynine can constitute up to 38.7% in traditional and commercial kratom products [5, 11, 12]. 7-Hydroxymitragynine is produced by oxidation of mitragynine and is a minor constituent (< 0.01% in fresh leaves) found at concentrations of up to 2% in leaf extracts and commercial kratom products [13–14]; however, it is believed to be the major contributor to the known addictive potential of kratom given its activity as a potent μ-opioid receptor agonist [15–18]. In the U.S., commercially available, imported kratom products (in the format of capsules, dried leaves, powders, resins, and concentrated extracts) have variable concentrations of mitragynine (1.2 to 38.74%) and 7-hydroxymitragynine (0.01 to 0.75%) on a leaf dry weight basis [11, 19]. Other major and minor alkaloids found within leaves of kratom include paynantheine (0.3 to 12.8% of kratom leaf dry weight), speciogynine (0.1 to 5.3%), and mitraphylline, which act as a competitive antagonist of μ-opioid receptors and function as muscle relaxants, and speciociliatine (0.4-12.3%) and corynantheidine (0.1-1.2%), which act as opioid agonists and adrenergic receptor [6, 10, 20–21]. The overall effect following consumption of kratom leaves is complex due to the interplay and range of bioactive alkaloids present [22].

Despite kratom’s long history of use in Southeast Asia, information on factors that influence plant growth and alkaloidal synthesis are largely unavailable. Available research on kratom is largely focused on its leaf chemistry and its potential pharmacological applications. As interest in cultivation of kratom increases along with consumptive demand, formal kratom cultivation efforts will likely soon be established. Although kratom had long been cultivated in Thailand, it was classified as illegal in 1943, and thus planting, possession, sales, and use of kratom leaves were prohibited [23]. In 2019, the Thai government approved a bill legalizing kratom for medicinal applications while recreational use remains illegal [24]. This bill provides the first opportunity for legal cultivation of kratom in Thailand since passage of the Kratom Act of 1943 and the Narcotics Act of 1979. Similar to other agricultural production efforts, plant cultivation practices based upon empirical evidence will be necessary for consistent successful commercial cultivation of kratom.

Synthesis of phytoactive leaf alkaloids can occur in response to light intensity. Highest natural photosynthetic light intensity, or photosynthetic photon flux density (PPFD), occurs outdoors in full-sun conditions during summer in the northern hemisphere. Plants respond to high PPFD by upregulating or downregulating alkaloid synthesis dependent upon species and other environmental factors present. For example, camptothecin, an indole alkaloid in *Camptotheca acuminate* leaves, was reduced by 99% when plants were moved from full sun to heavy shade (27% full sun) [25]. Similarly, total alkaloid content in tubers of *Pinellia ternate* decreased 27% when plants were moved from full sun to heavy shade (15% full sun) [26]. Light intensity in combination with nutrient availability may collectively affect phytoactive alkaloid production in some plant species. Winters and Loustalot (1952) observed limited synthesis of alkaloids in roots of *Cinchona ledgeriana* seedlings when subjected to 30% of full sunlight across a range of nitrogen fertility regimes [27]. However, considerable alkaloid synthesis did occur when light intensity was high and availability of nitrogen was low.

Phytoactive alkaloid synthesis within leaves can be affected by light quality, especially ultraviolet (UV) and far-red light, although studies are limited. Light quality is defined as the spectral composition of wavelengths influential to plant growth and photosynthesis. Concentration of the alkaloids catharanthine and vindoline in cell suspension culture of *Catharanthus roseu*s was promoted 3- and 118-fold, respectively, after being exposed to UV-B irradiation for a duration of 48 h as compared to plants not exposed to UV-B [28]. In addition to the influence of UV light, red and far-red light can influence plant secondary metabolism responsible for synthesis of leaf alkaloids. A low red to far-red light ratio, which occurs naturally in high shade or dense canopy conditions, results in a low phytochrome stationary state that can cause induction of shade avoidance responses, such as internode elongation and increase of leaf area and leaf chlorophyll concentration to optimize photosynthesis efficiency in the presence of competing vegetation [29–31]. However, the impact of light quality on alkaloid synthesis is still relatively unclear and largely undocumented. Tso et al. (2008) observed that total alkaloid content in tobacco (*Nicotiana tabacum*) tended to be higher in plants that were subjected to end-of-day red than far-red radiation; however, differences were not significant [32]. Although wild populations of kratom have been documented in the dense equatorial rain forests of Thailand and Malaysia, the influence of light on growth and alkaloid synthesis is undocumented and is vital to future commercial kratom cultivation efforts.

The sun is the major light source used in the cultivation of plants, even within most greenhouses where structural components may reduce or modify light intensity and quality. Quality of natural sunlight is strongly influenced by solar declination, atmospheric gases, and suspended particles that attenuate transmission of light. Light quality within greenhouses differs from outdoor, full sun conditions due to the absorptive nature of greenhouse structural materials, coverings, and any light-altering paints [33]. Furthermore, supplemental shade materials can be used to reduce light intensity and/or alter spectral quality when compared to outdoor, full sun conditions. Maximum daily light integral (DLI) outdoors within the state of Florida (U.S.) is approximately 45 mol·m^−2^·d^−1^ on a cloudless day in the summer, while the minimum may be less than 20 mol·m^−2^·d^−1^ during a short winter day in northern Florida [34]. More recently, we conducted a new data analysis from the NASA POWER database and discovered that the average DLI from 2015 to 2020 for Central Florida, Southern Thailand, Northern Malaysia, and Indonesia was 17.2, 17.3, 17.3, and 17.0 μmol·m^-2^·d^-1^, respectively. Although DLI in Central Florida is more variable throughout the year compared to Southeast Asia, mean DLI is similar. Depending upon glazing materials and structural components used, light transmission and intensity within a typical greenhouse is approximately 35-50% less than what is measured outdoors in full sun [35].

Although wild populations of kratom are found in the dense understory of equatorial rain forests, open-canopy commercial farming has recently been established in Indonesia in response to high export demands [36]. The influence of high light on plant growth and alkaloid content under open canopy production conditions, however, is undocumented. To determine if light-induced environmental factors can modify or influence synthesis of kratom leaf alkaloids, a preliminary investigation was conducted where kratom trees cultivated in a greenhouse were sampled to quantify leaf alkaloid content, moved to an outdoor, full-sun environment, and then sampled again two weeks later [37]. Concentrations of several leaf alkaloids increased in response to the change in cultivation environment, thus suggesting that an increase in PPFD, an increase in air temperature, a change in light quality, or a combination of these environmental factors were influential to synthesis of alkaloids within leaves of kratom. Given these preliminary findings, coupled with increased demand for kratom and a lack of foundational information regarding its cultivation, empirically derived information is needed by growers and producers to assist in attainment of biomass yield and alkaloid production goals. To directly address this need, research was conducted to examine the influence of light on kratom: 1) growth, leaf area, and biomass, 2) leaf chlorophyll content and its estimation through a rapid, non-destructive technique, and 3) the concentration of nine leaf alkaloids. Research results provide an expanded foundational knowledge of kratom and its response to varying light environments. This information will be helpful to the newly emerging commercial kratom cultivation industry where optimization of operations will be key to efficient and predictable production.

## Materials and Methods

### Plant Materials

Vegetative propagules, or cuttings, were taken from a single mother stock kratom plant, treated with 1000 mg·L^-1^ indole-3-butyric acid rooting hormone (Hormodin 1, OHP Inc., Mainland, PA, United States), and then placed within rockwool cubes to develop roots. Once roots had visually emerged from the rockwool cubes, the propagules were transplanted and cultivated in 0.7 L and 11.4 L containers as described by Zhang et al. (2020) [4]. Osmocote Plus 15-9-12 slow-release fertilizer (Scotts, Marysville, OH, United States) was applied at 74 g per container as per the manufacturer’s recommendations to provide sufficient nutrient availability throughout the duration of the experiment.

### Experiment Treatments

Sixty plants (n=60) were randomly assigned to one of three diverse light treatments to include: outdoor full sun, unshaded within a greenhouse (GH unshaded), and shaded within a greenhouse (GH shaded). Plants within the outdoor full sun treatment were placed under full sunlight outside of the greenhouse. Plants within the GH unshaded treatment were placed onto a bench within an enclosed greenhouse to receive ambient solar radiation. Lastly, plants within the GH shaded treatment were placed onto a bench inside of the greenhouse under shade cloth. Polycarbonate glazing materials reduced the daily light integral within the greenhouse (GH shaded) by approximately 60% compared to outdoor full sunlight. A knitted shade cloth (DeWitt, Sikeston, MO) installed approximately 2 m above a greenhouse bench reduced light by another 40% (~25% of full sun) to create conditions for the GH unshaded treatment. Environmental conditions within treatment areas were measured and adjusted to ensure limited variability existed. All plants were grown under natural day length regardless of treatment.

### Environmental Conditions

Plant irrigation schedule and greenhouse environment was monitored as described by Zhang et al. (2020) [4]. Outdoor environmental conditions were recorded every 15 min by the onsite Florida Automated Weather Network station. Average temperature within the greenhouse and the field were relatively similar throughout the experiment, with a mean of 27.7 to 28.1 ℃ in September, 24.3 to 25.0 ℃ in October, 19.9 to 21.7 ℃ in November, and 17.4 to 18.9 ℃ in December 2018.

Data collection protocol for this research was described previously in detail by Zhang et al. (2020) [4]. Briefly, plant height, width, trunk diameter, and SPAD value (an index of relative chlorophyll concentration) of mature leaves were collected monthly beginning September 10, 2018. Total leaf number and area, average leaf size, and total leaf dry mass was recorded at termination of the experiment on December 20, 2018. Specific leaf area was calculated by total leaf area and leaf dry mass. Quantification of leaf alkaloids and chlorophyll concentration was conducted monthly using the methods described in Zhang et al. (2020) [4]. In brief, leaf chlorophyll content was extracted and measured with a UV-Visible Spectrophotonmeter from three random plants within each treatment once every four weeks. A multiple reaction mode (MRM) based UPLC-MS/MS method in positive ionization was implemented for the quantification of nine kratom alkaloids on Acquity Class I UPLC coupled with Waters Xevo TQ-S Micro triple quadrupole mass spectrometer. UPLC method, compound and source parameters were the same as reported previously [4].

### Experiment Design and Data Analysis

The experiment was conducted using a complete randomized design with three treatments and 20 replicates. Each plant was considered as an experimental unit and an individual leaf sample was considered a subsample within the experimental unit. Statistical analysis was conducted using a restricted maximum likelihood mixed model analysis in JMP^®^ Pro 13 (SAS Institute, Inc., Cary, NC, United States) and SAS (SAS Institue, Inc., Cary, NC, United States). Post-hoc mean separation tests were performed using Tukey’s honest significant difference test by lighting treatment with treatment combination replicates (n=20) defined as the random error term. Statistical tests were considered significant if *P* ≤ 0.05.

## Results

### Plant Growth Indicators

Height of plants cultivated within the greenhouse was greater than those grown outdoors under full sunlight, with the tallest plants resulting from the shade cloth treatment (Fig. 1). Plant height was increased by 93 and 114% in response to being cultivated within the greenhouse under unshaded and shaded conditions, respectively, compared to plants cultivated outdoors under full sunlight. Similar to plant height, plants cultivated within the greenhouse were wider; plant width was between 53 and 57% greater than those cultivated in the field. Despite differences in height and width in response to imposed light treatments, trunk caliper growth of plants was similar among all treatments.

**Fig 1.**
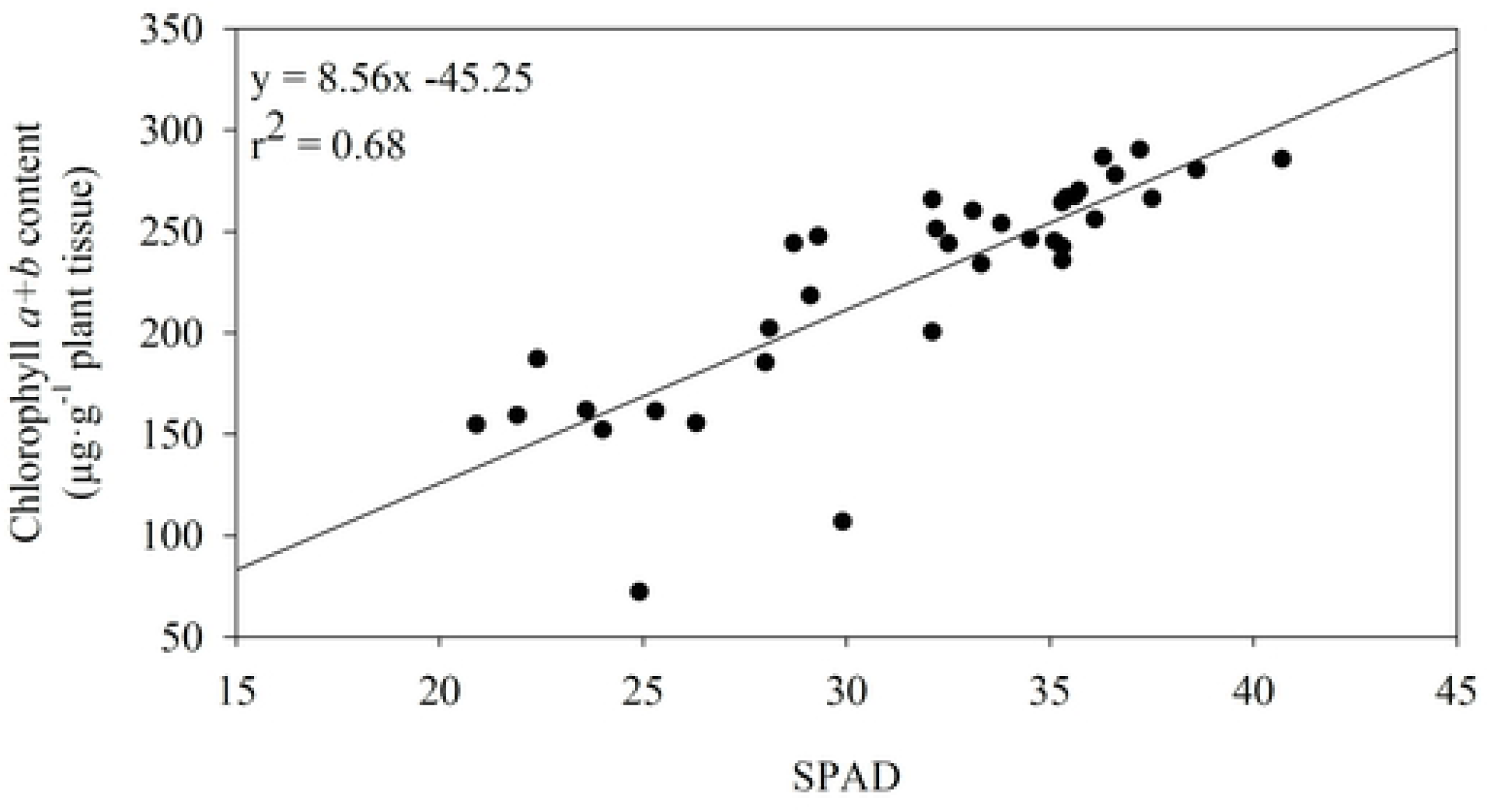
Plant growth indicators of kratom cultivated under varying radiance. GH = greenhouse. Leaf number included leaves ≥ 2 cm. Data were pooled from twenty replicates for height, width, and trunk caliper growth and four replicates for leaf number, total leaf area and average leaf size. Means sharing the same letter are not statistically different by Tukey’s honest significant difference test at *P ≤ 0.05*. NS = not significant.

Total leaf area and the number of leaves on plants grown inside the greenhouse (shaded and unshaded) were statistically similar and were 118 to 160% and 54 to 80% greater, respectively, than plants grown outdoors under full sun (Fig. 1). Average leaf area was similar between plants grown outdoors under full sun and those cultivated in the greenhouse without shade cloth. However, plants grown under shade cloth in the greenhouse had between 41 and 69% greater average leaf area than those cultivated in the unshaded greenhouse and under full sun, respectively. Total leaf dry mass trends were similar to that observed for total leaf area, plant width, and height with unshaded and shaded greenhouse cultivated kratom having 89 to 91% greater total leaf dry mass than those cultivated outdoors under full sun (Fig. 2). Additionally, specific leaf area increased with the decrease of light received, at 16 and 39% higher under the greenhouse unshaded and shaded conditions compared to full sun outdoors.

**Fig 2.**
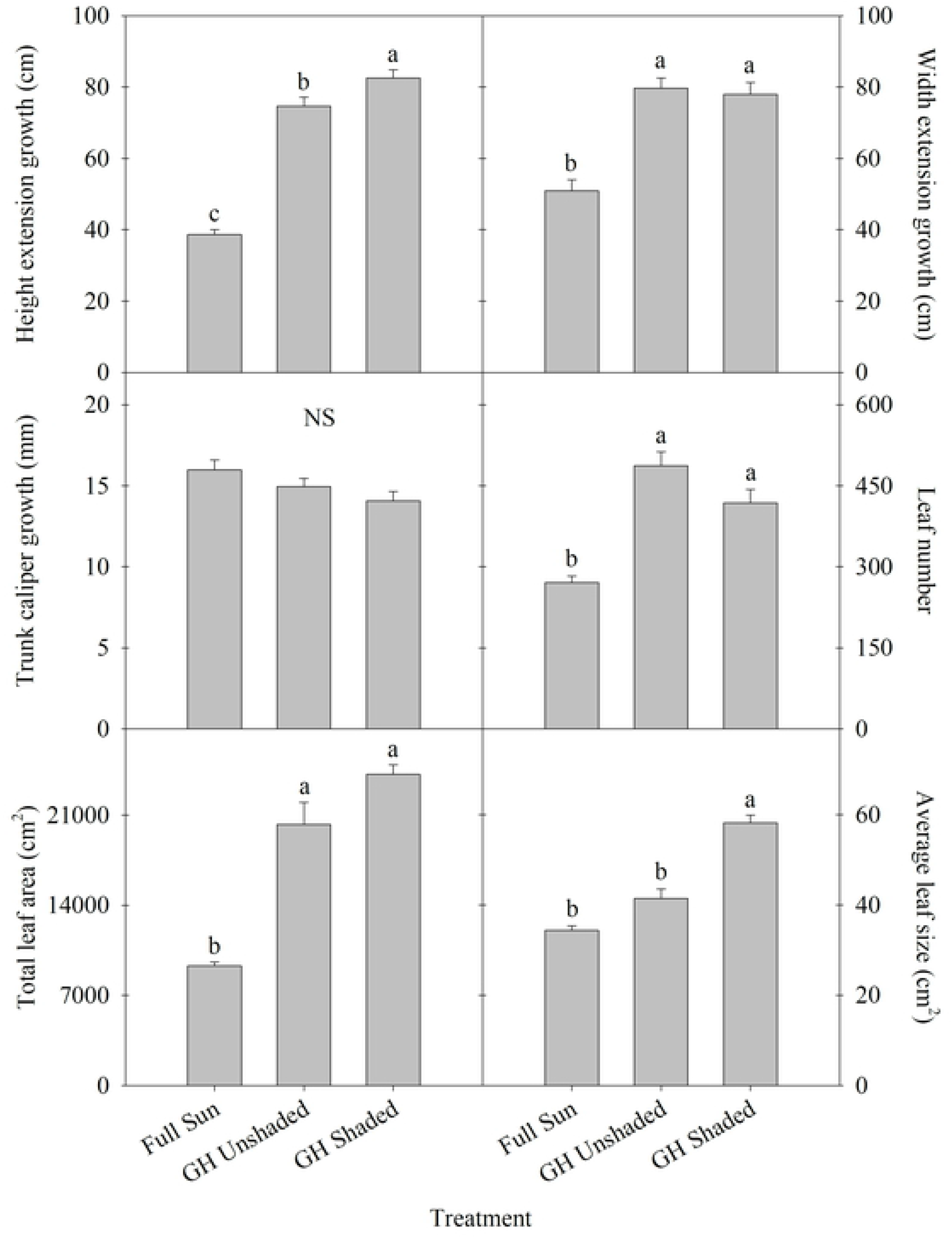
Total leaf dry mass and specific leaf area of kratom cultivated under varying radiance. GH = greenhouse. Data were pooled from four replicates and means sharing the same letter are not statistically different by Tukey’s honest significant difference test at *P ≤ 0.05*.

### SPAD and Chlorophyll Concentration

SPAD values were 22 to 31% greater among plants cultivated under shade cloth in the greenhouse than those grown in unshaded or under full sunlight in the field, respectively (Fig. 3). Chlorophyll concentration among treatments was similar at the beginning of the experiment and generally increased during the first month. Chlorophyll concentration of plants grown under full sun was 17% lower than those grown under shade cloth in the greenhouse, but both were not significantly different from plants grown under unshaded light in the greenhouse. At the end of the experiment, leaf chlorophyll concentration of plants grown under full sun was 17 to 23% less than those grown inside the greenhouse, but no differences were found between greenhouse treatments. Generally, chlorophyll a/b ratio was greatest when plants were grown under shaded conditions in the greenhouse, but no significant differences were found among treatments. Additionally, SPAD values correlated well (r^2^ = 0.68) with chlorophyll concentrations among trees between September and December 2018 (Fig. 4).

**Fig 3.**
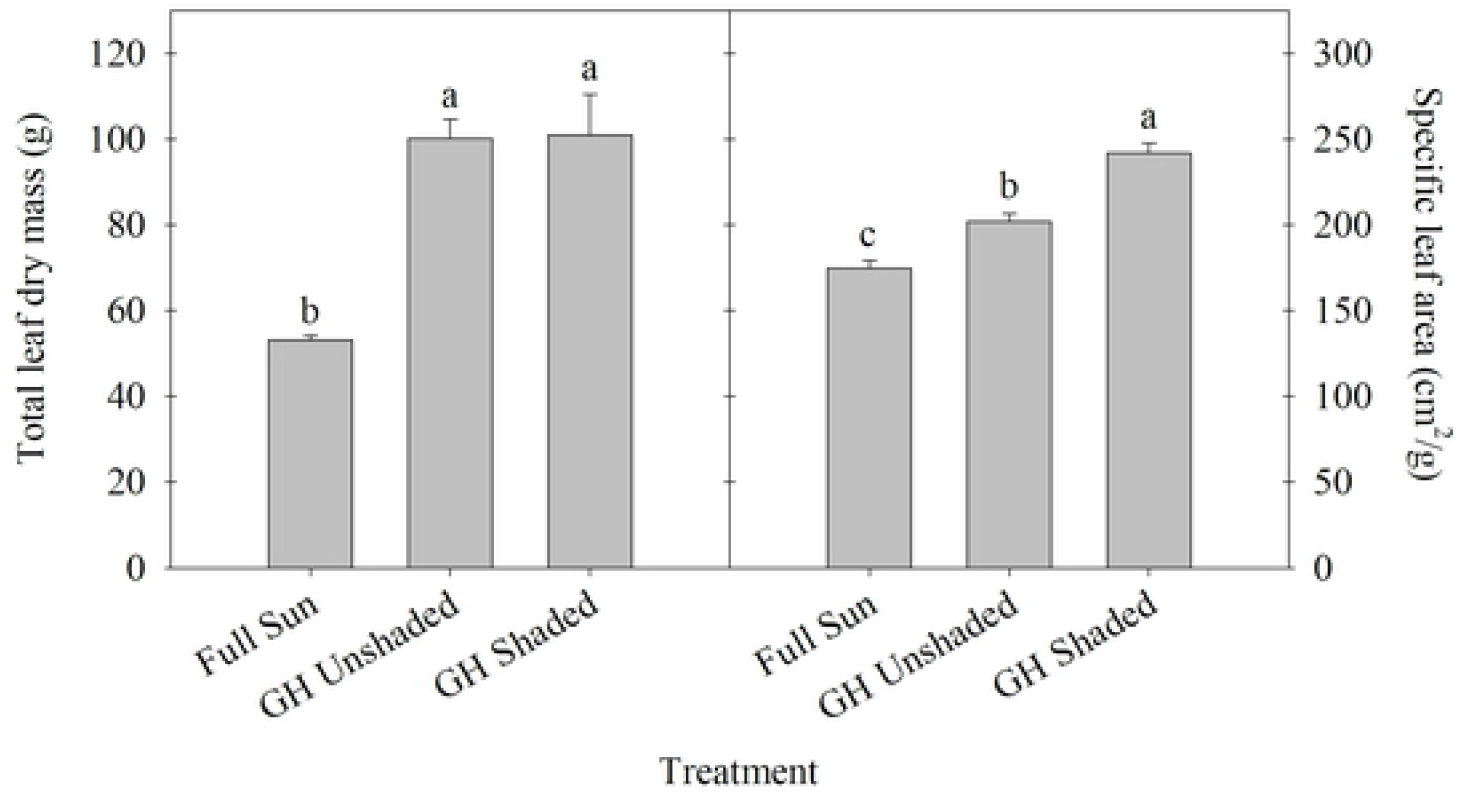
SPAD index values, chlorophyll concentration, and chlorophyll a/b ratio of kratom cultivated under varying radiance. GH = greenhouse. SPAD data were pooled from twelve replicates and chlorophyll *a+b* content and a/b ratio data were pooled from three replicates in each month. Means sharing the same letter are not statistically different by Tukey’s honest significant difference test at *P ≤ 0.05*.

**Fig 4.**
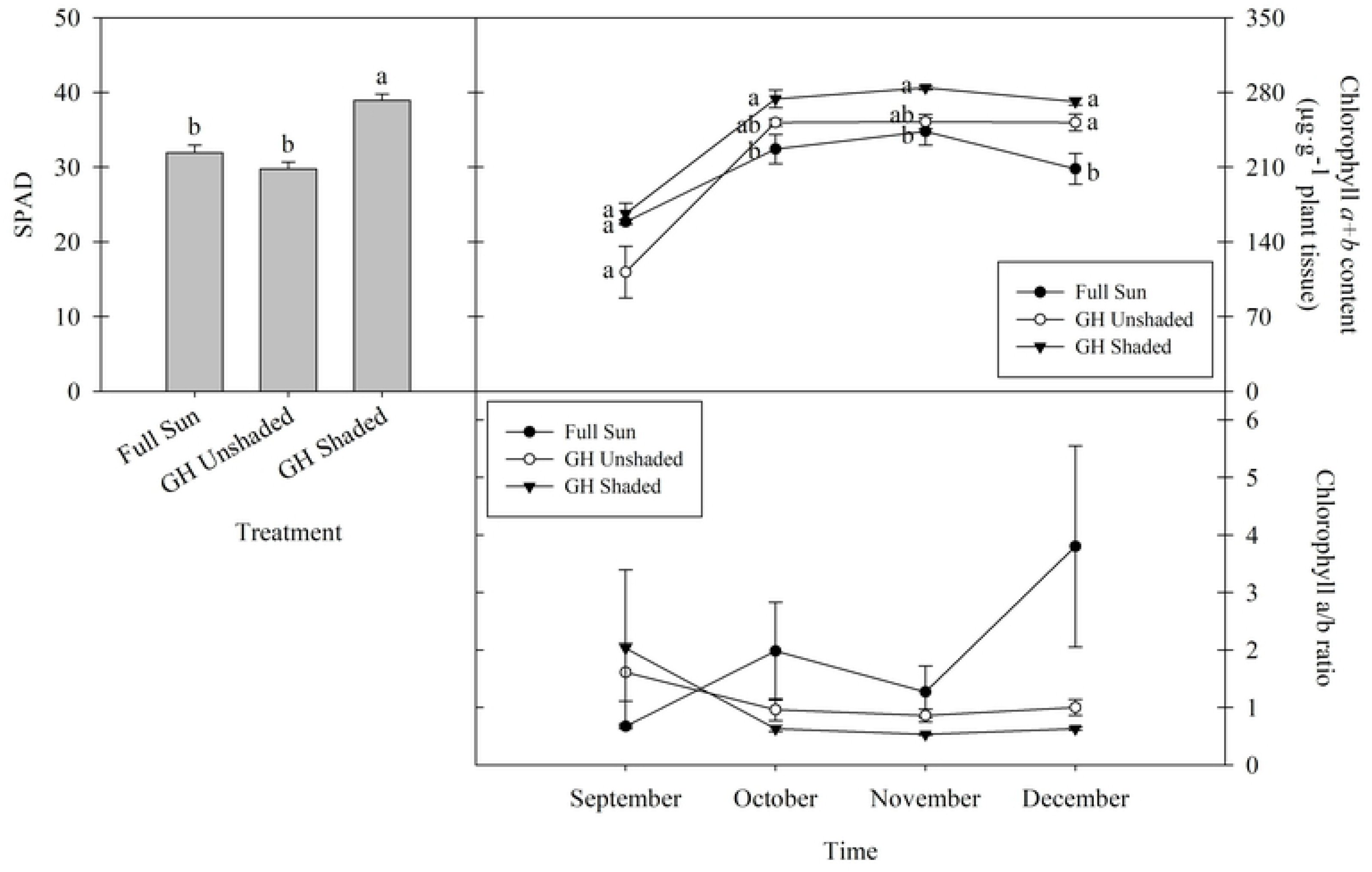
Correlation of SPAD and chlorophyll *a+b* content in kratom leaves. Plants were cultivated from September to December, 2018, under different radiation treatments. Data were pooled from 9 replicates each month.

### Alkaloid Concentration

7-hydroxymitragynine was not detected in any of our samples (Table 1). Mitragynine was detected in 53% of the samples; however, no significant differences among lighting treatments per leaf dry mass were observed (Table 1). On a per leaf dry mass basis, paynantheine concentrations in kratom grown under shade in the greenhouse were 40 and 27% greater than plants grown outdoors under full sun and in the greenhouse without shade, respectively. Similarly, corynoxine concentrations were approximately 2-fold greater in leaves of plants cultivated under shade in the greenhouse when compared to plants cultivated in full sun or in the greenhouse without shade. Despite these trends, differences in concentrations of speciogynine, speciociliatine, mitraphylline, corynantheidine, and isocorynantheidine in response to lighting treatment were not observed.

**Table 1.**
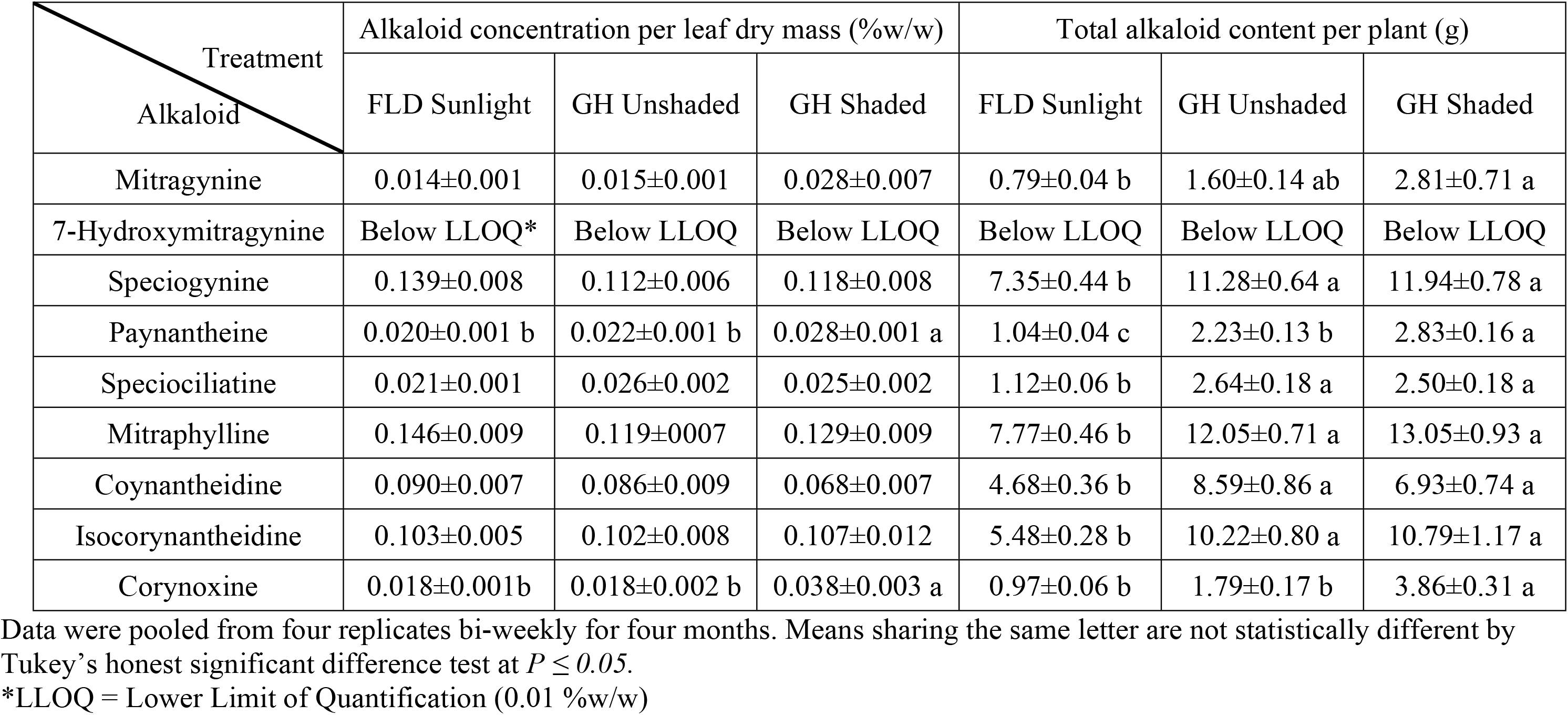
Phytoactive alkaloid concentration per leaf dry mass (±SE) and total phytoactive alkaloid content per plant (±SE) grown under field (FLD) sunlight or in greenhouse (GH) unshaded or shaded.

| Treatment<br>Alkaloid | Alkaloid concentration per leaf dry mass (%w/w) |  |  | Total alkaloid content per plant (g) |  |  |
| --- | --- | --- | --- | --- | --- | --- |
|  | FLD Sunlight | GH Unshaded | GH Shaded | FLD Sunlight | GH Unshaded | GH Shaded |
| Mitragynine | 0.014±0.001 | 0.015±0.001 | 0.028±0.007 | 0.79±0.04 b | 1.60±0.14 ab | 2.81±0.71 a |
| 7-Hydroxymitragynine | Below LLOQ* | Below LLOQ | Below LLOQ | Below LLOQ | Below LLOQ | Below LLOQ |
| Speciogynine | 0.139±0.008 | 0.112±0.006 | 0.118±0.008 | 7.35±0.44 b | 11.28±0.64 a | 11.94±0.78 a |
| Paynantheine | 0.020±0.001 b | 0.022±0.001 b | 0.028±0.001 a | 1.04±0.04 c | 2.23±0.13 b | 2.83±0.16 a |
| Speciociliatine | 0.021±0.001 | 0.026±0.002 | 0.025±0.002 | 1.12±0.06 b | 2.64±0.18 a | 2.50±0.18 a |
| Mitraphylline | 0.146±0.009 | 0.119±0.007 | 0.129±0.009 | 7.77±0.46 b | 12.05±0.71 a | 13.05±0.93 a |
| Coynantheidine | 0.090±0.007 | 0.086±0.009 | 0.068±0.007 | 4.68±0.36 b | 8.59±0.86 a | 6.93±0.74 a |
| Isocorynantheidine | 0.103±0.005 | 0.102±0.008 | 0.107±0.012 | 5.48±0.28 b | 10.22±0.80 a | 10.79±1.17 a |
| Corynoxine | 0.018±0.001b | 0.018±0.002 b | 0.038±0.003 a | 0.97±0.06 b | 1.79±0.17 b | 3.86±0.31 a |

Data were pooled from four replicates bi-weekly for four months. Means sharing the same letter are not statistically different by Tukey's honest significant difference test at $P \leq 0.05$ . \*LLOQ = Lower Limit of Quantification (0.01 %w/w)

On a per plant basis, total speciogynine, speciociliatine, mitraphylline, corynantheidine, and isocorynantheidine content of plants grown in the greenhouse, regardless of shading conditions, were approximately 1.6- to 2.4-fold greater compared to plants grown under full sun outdoors. Greenhouse shaded conditions drastically promoted total mitragynine synthesis (2.6-fold greater than plants cultivated outdoors under full sun). Similarly, total paynantheine content was 1.7- and 1.1-fold greater under greenhouse shaded conditions and unshaded conditions, respectively, than when cultivated outdoors under full sun.

## Discussion

Various species have shown to exhibit shade-acclimation response when subjected to shade or a low red to far-red light ratio environment to maximize sunlight interception [38–40]. In our study, kratom height, average leaf size, and total leaf dry mass were increased in response to unshaded and shaded conditions in the greenhouse compared to plants grown under full sun outdoors. Plants also developed significantly more specific leaf area and total chlorophyll content with a reduction in the chlorophyll a:b ratio, suggesting an optimization of light capture and a higher efficiency of light use in response to maximize photosynthesis and gain carbon under shaded conditions [41–42]. Similar findings have been observed in other related studies. For example, the height and leaf area of poinsettia (*Euphorbia pulcherrima*) were 55 to 75% and 111 to 155% greater when grown under 48 to 78% shading compared to 30% shading [43]. In addition, total chlorophyll content was highest under 92% of shade and dry weight was greatest under 48% of shade. In four different species of Pacific Northwest conifer seedlings, plant height was greatest and chlorophyll a was consistently higher under 75% of shade compared to no shade [44]. These shade-acclimation changes likely assist kratom in being more competitive in the dense, light-limited tropical forests in which they evolved and provide evidence of shade-acclimation response within this species. Given that leaf chlorophyll content was reliably estimated using SPAD meter values and fit a linear model developed in our study, cultivators of kratom may choose to reliably predict chlorophyll content using the rapid, nondestructive technique offered by the SPAD device.

Although originally believed to be byproducts of plant primary metabolic processes, phytoactive alkaloids are now better understood to be purposefully produced by plants to protect against herbivory and disease [9]. Relationships between environmental stimuli and the regulation of alkaloidal synthesis are complex and highly variable among plants and environments. In our study, paynantheine and corynoxine had the highest concentration per leaf dry mass under the most shaded lighting conditions. This is supported by Ralphs et al. (1998) where short-term shade stress, induced by 30% full sunlight for three days using shade cloth and 100% of full sunlight by covering leaves with aluminum foil, increased alkaloid concentration in tall larkspur (*Delphinium barbeyi*) by 36-38% and 11%, respectively, compared to plants grown in open sun [45]. Similarly, several plant alkaloids including vinblastine (*Catharanthus roseus*) and camptothecin (*Camptotheca acuminata*) have been reported in higher concentrations following exposure to low light conditions [25, 46]. However, contradictory relationships also exist within available related literature. For example, alkaloidal content in Kiasahan (*Tetracera scandens*) leaves was higher under full sunlight than under shade [47]. Similarly, Chen et al. (2017) reported an increase in total alkaloid content in *Pinellia ternate* when light was increased from 15 to 100% of full sunlight using shade nets [26]. This is not uncommon as alkaloid production involves several different metabolic pathways, although many of them are not fully understood [48]. Although thought to be an overly simplified relationship, our findings support the carbon/nutrient balance theory where carbon stress due to limitation of light and a resulting reduction in photosynthesis increase nitrogen-containing defense compounds, such as alkaloids, in shade-tolerant species [49]. Additionally, a high specific leaf area for plants growing in the shade can make their leaves more sensitive to mechanical stress and herbivory, thus an increasing level in alkaloids may assist with the defense mechanism for survival in deep shade [41, 48, 50].

Cultivating kratom under shade cloth within a greenhouse maximized the concentration of both paynantheine and corynoxine. In addition, this same production condition maximized plant height, leaf area, and leaf size. Given this, total calculated yield of each alkaloid quantified in our study was greatest among shaded plants given the shade acclimation response of greater leaf mass and a larger leaf size (Table 1). An evolutionary adaptation to low-light environments is likely given the higher plant performance observed when kratom was cultivated in conditions similar to those that occur in the dense, shaded understory of equatorial rainforests. Given that paynantheine and corynantheidine act as muscle relaxers and opioid agonists, cultivators of kratom could produce plants under similar production conditions to maximize production and use of kratom for these intended applications. Despite the range of lighting conditions imposed in this study, only low levels of mitragynine were observed. 7-hydroxymitragynine was not detected in any leaf samples, reinforcing the opinion that this alkaloid is produced from mitragynine as a post harvest artifact [5]. Low levels of mitragynine, coupled with a lack of 7-hydroxymitragynine, suggests low abuse liability potential when compared to previously examined imported commercial kratom products.

When conducting horticultural investigations, differences in irradiance usually accompany a differential in environmental temperature; however, environmental temperature information is often not discussed, reported, or otherwise accounted for in available literature. In our study, greenhouse and field temperatures were managed so they remained similar throughout the experiment and thus variable temperatures among imposed treatments were eliminated as a potential confounding variable. Research examining synthesis of leaf alkaloids in response to different temperatures under similar light intensities in the controlled environments is warranted and remains needed, however.

In addition to being influenced by light intensity and temperature, synthesis of phytoactive alkaloids may be influenced by light quality. Indole alkaloid concentrations have been observed to vary in response to UV-B irradiation exposure in a number of medicinal plants including *Clematis terniflora*, *Withania somnifera*, *Coleus forskohlii*, *Zanthoxylum bungeanum*, and *Coleus aromaticus* [51]. Ultraviolet light transmission is often 20 to 80% lower within greenhouses than outdoors in full sun, dependent upon glazing materials used in the construction and design of the greenhouse [52]. Surprisingly, in our study alkaloid concentrations were not different between plants cultivated under full sunlight outdoors and those grown within the greenhouse without supplemental shading material. In our preliminary study, an increase of mitragynine was observed after plants were moved from within the greenhouse to outdoors; however, the increase was not significant [37]. Given that this preliminary study was only exploratory, plant number and experiment duration were limited, and the environment was not strictly controlled, we believe that this observation may have been caused by plant individual differences and not a light treatment effect. The preliminary study also relied upon a different analytical method to quantify alkaloid concentrations, thus confounding accurate comparisons between studies. Alternatively, differences may have been due to increased aging of greenhouse materials and degradation of the UV stabilizer found in its roof material. Thus, more UV radiation entered the greenhouse in this study and created little to no UV difference compared to the field [53]. Regardless, significant differences in the concentration of the alkaloids paynantheine and corynoxine were observed among plants subjected to shaded and unshaded light conditions within the greenhouse. In addition to reducing light intensity, shade cloth modifies light quality by causing a shift in the red to far-red light ratio. Together, results suggested that a change of light intensity, a change of light quality, or a combination of both may result in the alteration of leaf phytoactive alkaloids in kratom, particularly in the case of corynoxine where concentrations differed approximately 2-fold in response to lighting treatments.

## Conclusion

Given recent increased interest in the cultivation and application of kratom, foundational research examining the influence of light on kratom growth and alkaloid synthesis was conducted. Although light significantly influenced plant growth, it did not influence the synthesis of most leaf alkaloids. Only paynantheine and corynoxine concentrations, per leaf dry mass, increased under shade conditions when cultivated within a greenhouse. Despite minimal influence on the synthesis of leaf alkaloids, greenhouse lighting conditions drastically increased total leaf dry mass. Total alkaloid yield per plant, as a result, was maximized under low-light conditions. Given these findings, production efforts where low-light conditions can be implemented would likely maximize plant biomass and total alkaloid leaf concentrations.

## Acknowledgement

We gratefully acknowledge Kelly and Liz Dunn along with Darren Frankle for plant material and donations to the University of Florida Foundation for supporting the project. In addition, we gratefully acknowledge Scott Acker for his support of our kratom research efforts. We also thank Dr. Heather Enloe for her assistance with the manuscript.

## Author Contributions

**Conceptualization:** MZ, BJP.

**Data curation:** MZ, AS, FL.

**Formal analysis:** MZ.

**Funding acquisition:** BJP, RK, CRM.

**Investigation:** MZ, AS, FL.

**Methodology:** AS, FL, BA, CRM

**Supervision:** BJP, CRM.

**Visualization:** MZ.

**Writing – original draft:** MZ.

**Writing – review & editing:** MZ, AS, FL, RK, CRM, BJP.

## Notes

### Competing Interest Statement

The authors have declared no competing interest.

